# Differences between human and rodent nitric oxide production dictate susceptibility to tick-borne *Rickettsia*

**DOI:** 10.1101/2025.06.27.661835

**Authors:** Anh Phuong Luu, Alejandro Amando Guzman, Alexis Bouin, Nir Drayman, Thomas P. Burke

## Abstract

Arthropod-borne pathogens cause serious human infections, yet they only cause limited disease in rodent reservoirs. Wild type mice resist infection by tick-borne *Rickettsia parkeri,* which causes spotted fever in humans, and it remains unclear why humans are vulnerable. Here, we report that whereas mouse type I interferon (IFN-I) or interferon-γ (IFN-γ) dramatically restrict *R. parkeri* in macrophages, human interferons do not. Differential RNA-seq revealed a significant induction of nitric oxide synthase 2 (*Nos2,* encoding inducible nitric oxide synthase, iNOS) in infected mouse but not human macrophages upon interferon treatment. Chemical iNOS inhibition or *Nos2* deletion restored IFN-γ-mediated restriction in mouse cells. Human cells treated with cytokine cocktails or with iNOS cofactors and substrates were still unable to restrict *R. parkeri*. *In vivo*, whereas wild type mice restricted *R. parkeri*, infected *Nos2^-/-^*mice developed mild skin eschars, recapitulating a key human disease manifestation. Together, our findings suggest that there is a threshold of NO production required to restrict *R. parkeri,* which mouse cells reach but human cells do not, and this is a key explanation for why humans develop tick-borne rickettsial diseases while rodents can be tolerant, asymptomatic reservoirs. Differences in NO abundance may provide an evolutionary explanation for human susceptibility to pathogens that propagate themselves in rodent reservoirs.

## Introduction

Zoonotic pathogens often circulate in wildlife reservoirs, where they can persist without causing significant harm to the host. However, when spilling over to humans, many cause severe morbidity and mortality. For example, the Lyme disease agent *Borrelia burgdorferi* spreads horizontally from tick to tick in deer mice (Braks et al., 2016; Springer et al., 2021). In the case of many viruses, such as Dengue, Zika, and West Nile, humans serve as amplifying hosts for mosquitoes; yet, rodents often exhibit higher resistance to infection (Blahove & Carter, 2021; Lazear et al., 2016; Vandegrift et al., 2020). In the case of tick-borne *Rickettsia* species, humans worldwide suffer from severe spotted fever disease, while animals, which can propagate the bacteria in the wild, show much greater resistance (Dantas-Torres, 2007; Grasperge, Reif, et al., 2012; Peniche-Lara et al., 2015). The molecular explanations for why humans develop severe rickettsial disease, whereas rodents do not, are unclear. Elucidating these mechanisms is critical for better understanding human disease, elucidating how tolerant animals in the environment harbor pathogens, and for developing novel therapeutics to treat infection.

Spotted fever group (SFG) *Rickettsia* species are obligate intracellular, Gram-negative tick-borne bacteria, and they are transmitted to hosts during blood-feeding (Parola et al., 2005). The maintenance cycle of many pathogenic *Rickettsia* relies on arthropod vectors, including ticks, and amplifying hosts such as rodents and dogs (Alvarez-Londoño et al., 2025; Azad & Beard, 1998; Helminiak et al., 2022). The bacteria can also undergo transovarial transmission from adult ticks to ova that hatch into infected larval offspring (Kloc et al., 2024; Moraes-Filho et al., 2018). Humans, in contrast, are accidental or dead-end hosts that develop debilitating and sometimes fatal diseases (Alvarez-Londoño et al., 2025; Kim, 2022; Parola et al., 2013). Among SFG *Rickettsia* family members, *R. parkeri*, found in the Americas, causes a non-lethal eschar-associated rickettsiosis. Upon infection, SFG *Rickettsia* is initially distributed in the skin and regionally via lymphatics and then disseminated through the blood, where the more pathogenic species target vascular endothelial cells (Teng & Chatham, 2015; Walker, 1989). Rickettsial infection of the endothelium can cause a characteristic fever, headache, and myalgia. *R. parkeri* targets mononuclear cells, including macrophages, and causes a necrotic skin lesion at the site of a tick bite called an eschar, which is a hallmark disease manifestation of *R. parkeri* rickettsiosis and some other arthropod-borne diseases such as scrub typhus (C. D. Paddock et al., 2008; Paddock et al., 2004; Scott et al., 2022). Despite these serious manifestations in humans, wild-type mice are highly resistant to infection (Grasperge, Reif, et al., 2012), and we are pursuing the hypothesis that the molecular explanation for human susceptibility versus rodent resistance is dictated by innate immunity.

Innate immunity employs pattern recognition receptors (PRRs) to detect conserved bacterial components (Mogensen Trine, 2009). Once activated, PRRs trigger a series of signaling cascades, leading to the production of antimicrobial cytokines, including type I interferon (IFN-I) (Gongora & Mechti, 1999). Murine IFN-I or interferon-γ (IFN-γ) efficiently controls *R. parkeri* infection *in vitro* (Burke et al., 2020). *In vivo*, single mutant mice lacking either receptor for IFN-I (IFNAR) or IFN-γ (IFNGR) resist infection, yet double mutant *Ifnar^-/-^Ifngr^-/-^* mice develop necrotic skin lesions (Burke et al., 2021), resembling those seen in humans infected with *R. parkeri* (Cragun et al., 2010; Kaskas et al., 2014; C. D. Paddock et al., 2008). As *Rag2^-/-^* mice are highly resistant to *R. parkeri* (Burke et al., 2020), this reveals innate and not adaptive immunity is key for protection against acute infection. Similarly, the mite-borne pathogen *Orientia tsutsugamushi* also causes limited disease in WT mice but eschar-associated disease in mice lacking interferon signaling (Liang et al., 2024). The exact mechanisms of this restriction by interferons are unclear. IFN-I or IFN-γ activates the Janus kinase signal transducer and activator of transcription (JAK-STAT) signaling pathway (Platanias, 2005), inducing the expression of hundreds of antimicrobial host effector proteins called interferon-stimulated genes (ISGs) that target and restrict pathogens (Schneider et al., 2014). Many ISGs have been identified to be antimicrobial toward viruses and vacuolar bacteria, but only a few such as guanylate binding protein 2 (GBP2) and inducible nitric oxide synthase (iNOS) have been described to target obligate cytosolic bacteria (Burke et al., 2020; Fitzsimmons et al., 2021). This provides the premise for our hypothesis that mouse ISGs restrict *R. parkeri*, but humans lack or carry malfunctional anti-rickettsial ISGs.

iNOS is encoded by the *Nos2* gene and is upregulated downstream of IFNs to produce nitric oxide (NO), a diffusible radical gas (Mehta et al., 2012; Morikawa et al., 2000; Tötemeyer et al., 2006). NO has antimicrobial activities against multiple pathogens such as viruses, bacteria, fungi and protozoa (Bogdan, 2001b; MacMicking et al., 1997; Zhao et al., 2024). Together with NADPH oxidase-mediated reactive oxygen species (ROS), NO creates a variety of secondary reactive nitrogen species (RNS), such as peroxynitrite (ONOO^−^) and dinitrogen trioxide (N_2_O_3_), that attack the proteins, lipids, and DNA of invading pathogens (Fang, 2004; Wang & Ruby, 2011). While NO is produced in human macrophages, the amount produced is approximately >20-fold less than in mouse and rat macrophages *in vivo* and *in vitro* (Azenabor et al., 2009; Blond et al., 2000; Bogdan, 2001a, 2001b; Jiang et al., 2023; Panaro et al., 2003). While the differences between human and mouse iNOS have been investigated as perhaps an undesired consequence of working with mouse models, the biological consequences of this difference and its evolutionary role in arthropod-borne disease has not been appreciated.

To determine the molecular explanation for why humans develop rickettsial diseases while animal reservoirs are resistant, we performed RNA-seq analyses between IFN-treated mouse and human macrophages infected with *R. parkeri*. *Nos2* was the most abundantly increased gene in mouse cells but was not upregulated in human cells. Inhibiting iNOS in IFN-γ-treated mouse cells abolished the antimicrobial effect, whereas combinations of cytokine cocktails or supplementing iNOS substrates and cofactors did not restrict *R. parkeri* in human cells. *In vivo*, *Nos2^-/-^* mice developed mild eschars with no lethality, providing the closest animal model yet to mimic *R. parkeri* rickettsiosis in mice. Together, our findings may suggest that reduced NO production is a key explanation to why humans, which rarely encounter such pathogens, develop tick-borne rickettsial diseases while rodents, which frequently interface with ticks in nature, maintained resistance.

## Results

### Primary mouse but not human macrophages control *R. parkeri* upon IFN-I or IFN-γ treatment

We sought to decipher the mechanisms behind why humans develop rickettsioses while mice do not. As *R. parkeri* resides within macrophages in mice and is restricted by mouse interferons (Burke et al., 2021), and since *R. parkeri* resides in macrophages in human skin biopsies (Herrick et al., 2016; Christopher D. Paddock et al., 2008), we hypothesized that human and mouse interferons had differing abilities to restrict *R. parkeri* in macrophages. To test this, we generated mouse bone marrow-derived macrophages (BMDMs), infected them with *R. parkeri*, treated the cells with recombinant mouse IFN-I or IFN-γ 10 minutes postinfection (mpi), and measure plaque forming units (PFUs) over time. Aligning to our previous findings, mouse IFN-I or IFN-γ restricted *R. parkeri* in BMDMs (**Fig. 1A**). To determine whether a similar degree of restriction occurred in human macrophages, we obtained blood from healthy human donors, purified monocytes with CD14+ selection, and derived them into macrophages with either M-CSF or GM-CSF. These monocyte-derived macrophages (MDMs) were infected with *R. parkeri,* treated with human IFN-I or IFN-γ, and PFUs were quantified over time. To account for any potential differences in the abundance of IFNs that may be produced in each species *in vivo*, we treated human macrophages with 10-fold more IFN-α and IFN-γ than mouse macrophages. Strikingly, while mouse IFN-I and IFN-γ restricted *R. parkeri* 37-fold and >100,000-fold, respectively, treatment of human macrophages with excessive IFNs only restricted 4-fold and 6-fold, respectively (**Fig. 1B).** As a control, we measured killing of vesicular stomatitis virus (VSV) and found that VSV was strongly restricted by human IFN-I in MDMs (**Fig. 1C**), verifying that the recombinant human IFN-I was functional. To further confirm the functionality of both IFNs in this system, we performed quantitative PCR (qPCR) on MDMs to measure ISG upregulation. Both human IFNs induced the expression of known ISGs (**Fig. 1D**), confirming that the IFNs were functional, and that the cells were responsive. These data suggest that rodent but not human macrophages induce anti-rickettsial factors and control infection in response to IFNs.

**Figure 1:**
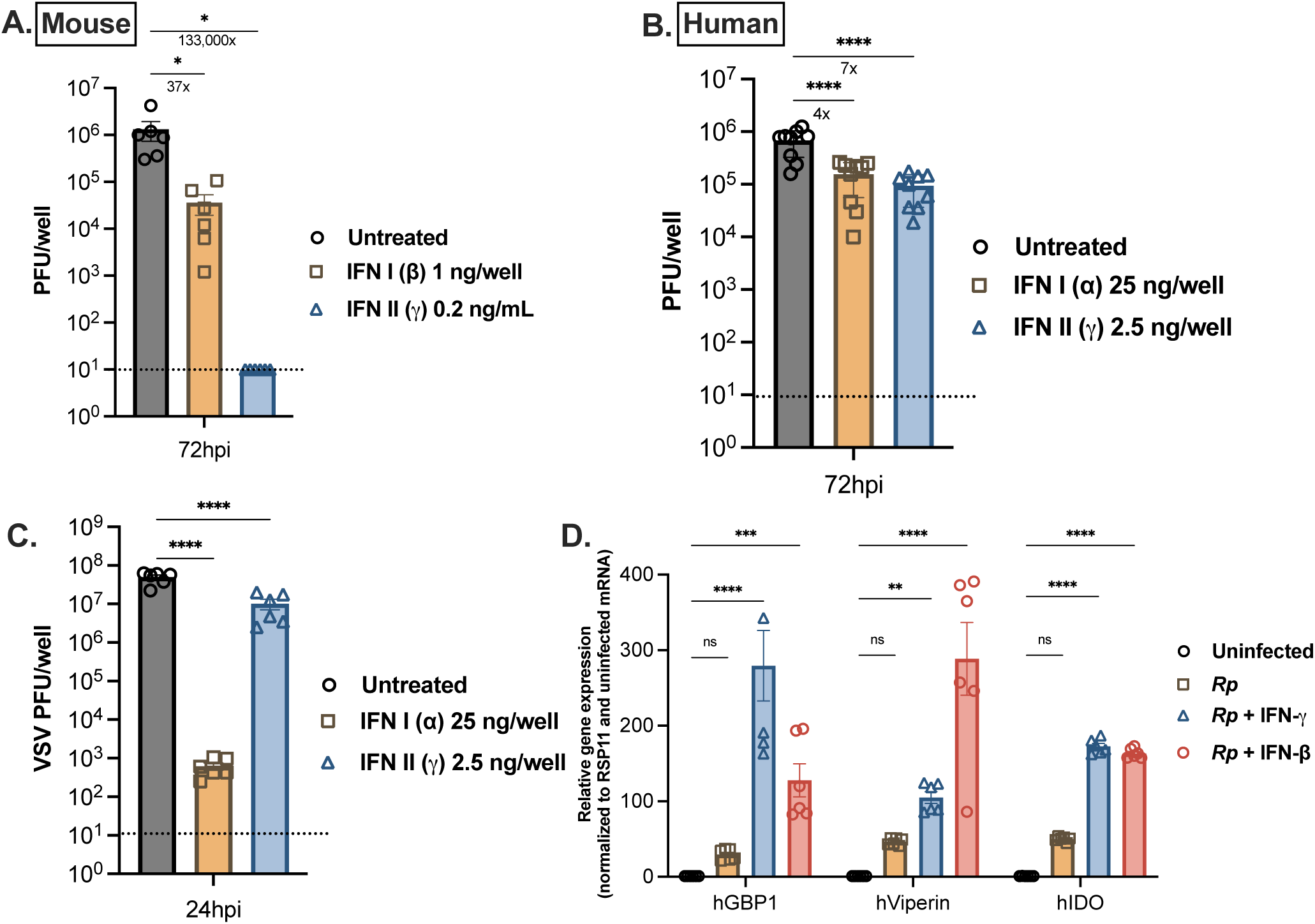
IFN-I and IFN-γ restrict *R. parkeri* in primary mouse but not human macrophages. **A.** *R. parkeri* abundance in mouse BMDMs at 72 hpi; MOI of 0.2. The indicated amounts of IFN-β and IFN-γ were added 10 min post-infection. Each data point is from a well of lysed infected cells from 6 independent experiments, where each experiment had 2 biological replicates. **B**. *R. parkeri* abundance in human MDMs at 72 hpi; MOI of 0.5. The indicated amounts of IFN-α and IFN-γ were added 10 min post-infection. Human MDMs were derived from 3 healthy donors. N=9, where each data point is from a well of lysed infected cells, and each experiment had 3 biological replicates. **C**. VSV titers from supernatants of infected primary human MDMs at 24 hpi, MOI of 1. The indicated amounts of IFN-α and IFN-γ were added 24 h before infection; human MDMs were derived from 3 healthy donors. N=6, where each data point is supernatant from a well of infected cells, and each experiment had 2 biological replicates**. D**. Gene expression in infected human MDMs at 12 hpi, MOI of 2, was quantified by qPCR. Data were normalized to RSP11 and then divided by uninfected to reveal fold change. Human MDMs were derived from 2 healthy donors. Each data point is RNA from a well of lysed infected cells, and each experiment had 3 biological replicates. Asterisks indicate statistically significant differences. All error bars indicate the standard error of the mean (SEM). Statistics used a one-way ANOVA with Dunnett’s test. *, p < 0.05; **, p < 0.01; ***, p < 0.001; ****, p < 0.0001.

### *Nos2* is the major anti-rickettsial gene upregulated by IFN-**y** in mouse macrophages

The specific ISG(s) responsible for restricting *R. parkeri* in mouse but not human macrophages were unknown. We conducted RNA-seq to compare the ISG profiles between infected mouse BMDMs and human MDMs in presence of IFNs. We chose a time point of 12 hpi because this was early enough to start to see restriction of *R. parkeri* (Burke et al., 2020), and we sought to minimize downstream indirect signaling that may occur later during infection. RNA was extracted and sequenced, and the transcriptomes of IFN-treated, infected cells was compared to that of untreated and uninfected cells using DESeq2. >3,000 genes were significantly upregulated (Log2 fold-change >2, adjusted p-value <0.05) in mouse macrophages compared to ∼700 genes in human macrophages (**Supplemental Table 1**). Among these, *Nos2* stood out as the most abundantly upregulated gene in mouse macrophages (>64,000-fold) but it was not significantly upregulated (unchanged) in human cells (**Fig. 2A**). *R. parkeri* infection itself (without IFN-γ) also led to a significant upregulation of *Nos2* in mouse but not human cells (>23,000-fold) and other genes (**Fig. 2A**). This high upregulation of *Nos2* by *Rp* alone in a scenario where the bacteria successfully replicate in these cells suggests that perhaps a very high amount of NO is required to restrict the pathogen. Given its known antimicrobial function against *R. rickettsii* and other pathogens, we hypothesized that iNOS was responsible for IFN-γ-mediated restriction of *R. parkeri* in mouse but not human cells.

**Figure 2:**
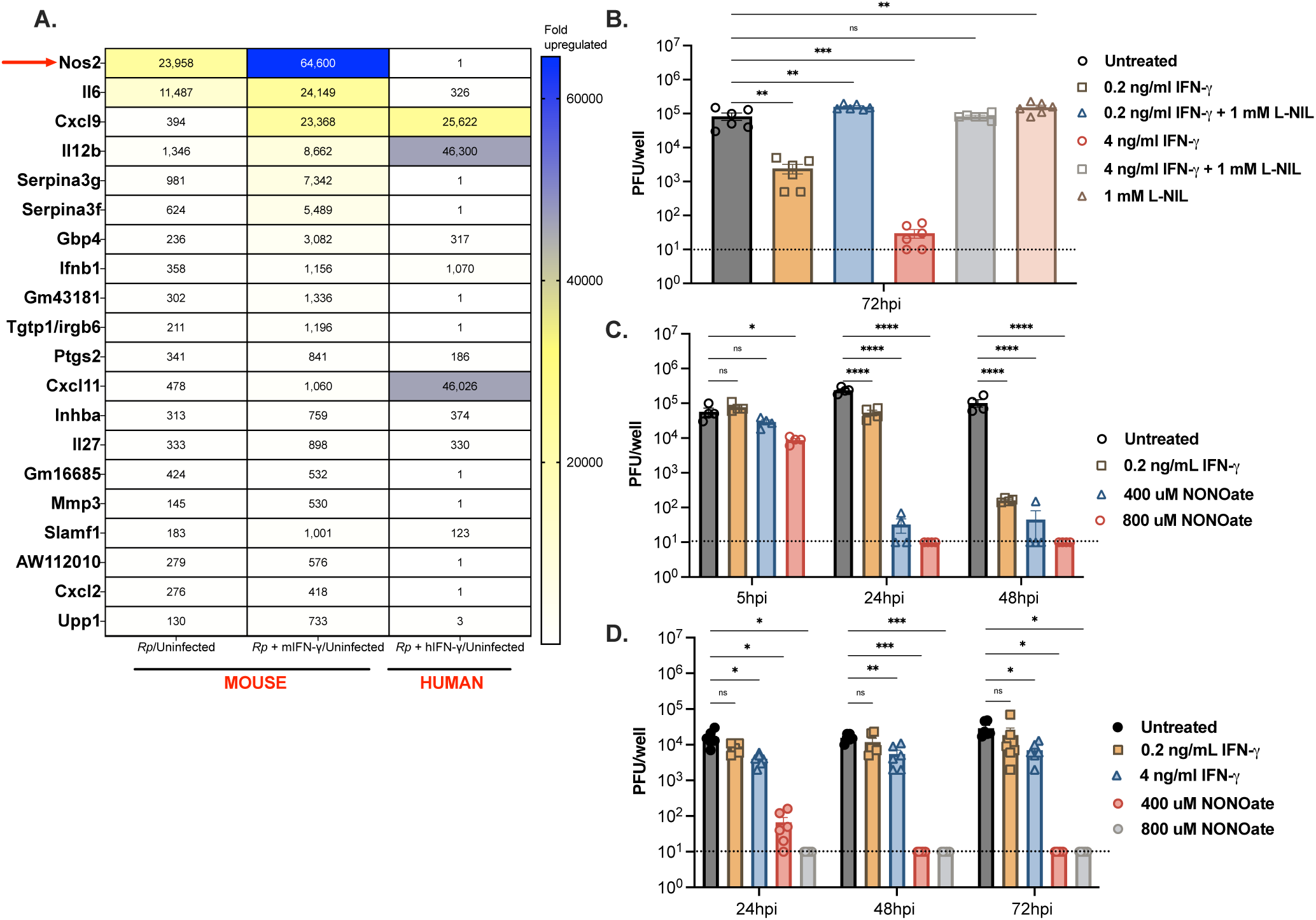
IFN-γ upregulates mouse but not human *Nos2*, which is essential for *R. parkeri* restriction in mouse cells. **A** Heatmap of upregulated transcripts in infected mouse BMDMs and human MDMs upon IFN-γ treatment at 12 hpi; MOI of 2. Data was normalized by dividing the averaged values of transcript counts of each sample over the averaged uninfected transcript counts; BMDMs and human MDMs were derived from 2 mice and 3 healthy donors, respectively. **B.** *R. parkeri* abundance in mouse BMDMs at 72 hpi; MOI of 1. The indicated amounts of IFN-γ and 1 mM L-NIL were added at 10 mpi. Data are the combination of 2 independent experiments with 3 biological replicates for each experiment. **C.** *R. parkeri* abundance in mouse BMDMs at the indicated times, MOI of 1. The indicated amount of IFN-γ was added at 10 mpi. NONOATE was added at 10 mpi and 24 hpi. Data are combined from 2 independent experiments with 2 biological replicates per condition in each experiment. **D**. *R. parkeri* abundance in mouse *Nos2*^-/-^ BMDMs at 24, 48, and 72 hpi; MOI of 0.2. The indicated amount of IFN-γ was added at 10 mpi. NONOATE was added at 10 mpi and 24 hpi. Data are combined from 3 independent experiments with 2 biological replicates each. Asterisks indicate statistically significant differences. Statistics used a one-way ANOVA for 2 B. Panels 2C and 2D used a two-way ANOVA with Dunnett’s multiple comparisons test: *, p < 0.05; **, p < 0.01; ***, p < 0.001; ****, p < 0.0001.

To determine the anti-rickettsial role of iNOS during IFN-γ treatment, IFN-γ-treated BMDMs infected with *R. parkeri* were treated L-NIL, a widely used and specific iNOS inhibitor (Moore et al., 1994). L-NIL completely reversed IFN-γ-mediated restriction of *R. parkeri* in BMDMs at 72 hpi upon treatment with either 0.2 ng/ml or 4 ng/ml IFN-γ (**Fig. 2B).** Similarly, BMDMs derived from *Nos2^-/-^* mice were unable to restrict *R. parkeri* upon IFN-γ treatment, verifying the results with L-NIL (**Fig. 2D).** This aligned with findings that IFN-γ induces abundant iNOS-produced NO in mouse cells to kill the SFG species *R. rickettsii* (Fitzsimmons et al., 2021) and suggested that iNOS is the major IFN-γ-regulated ISG responsible for killing *R. parkeri* in murine BMDMs.

It remained unclear if NO was sufficient to kill *R. parkeri* intracellularly. To determine if NO-mediated killing could be recapitulated in the absence of IFN-γ, the NO donor NONOATE was added to infected BMDMs and PFUs were measured over time. NONOate significantly inhibited *R. parkeri* even at early time points (24 hpi) in WT BMDMs (**Fig. 2C**) or in *Nos2*^-/-^ BMDMs (**Fig. 2D**). This demonstrates that NO produced by iNOS is the major IFN-γ-stimulated factor that restricts *R. parkeri* in mouse macrophages. Importantly, since *R. parkeri* infection itself upregulates iNOS but the bacteria still grow, this suggests that IFN-γ enables cells to meet a threshold of NO that can restrict *R. parkeri* intracellularly.

### iNOS induction and activation in human lung cells or macrophages does not strongly restrict *R. parkeri*

It was unclear if NO produced by iNOS could kill *R. parkeri* in human cells. Previous studies reported that combinations of IFN-γ, TNF-α, and IL-1β induced iNOS in A549 human lung epithelial cells (Barilli et al., 2023). We therefore sought to determine if cytokine cocktails were sufficient to induce *Nos2* in MDMs and A549s to restrict *R. parkeri*. In uninfected A549s, whereas IFN-γ induced iNOS 10-fold, IL-1β induced iNOS 100-fold. Combination treatments further induced iNOS, whereby IFN-γ plus IL-1β or the combination of IFN-γ, IL-1β, and TNF-α induced iNOS to ∼10,000-fold over uninfected controls (**Fig. 3A**). Despite the high upregulation of iNOS, *R. parkeri* abundance was only slightly reduced, even upon treating A549s with cytokine cocktails three times (**Fig. 3B**). In MDMs, IFN-γ, TNF-α and IL-1β treatment induced *Nos2* only 2-fold above infected cells (**Fig 3C**). This led us to conclude that iNOS expression alone was insufficient to elicit potent anti-rickettsial NO production and therefore we next investigated the role for iNOS substrates and cofactors.

**Figure 3:**
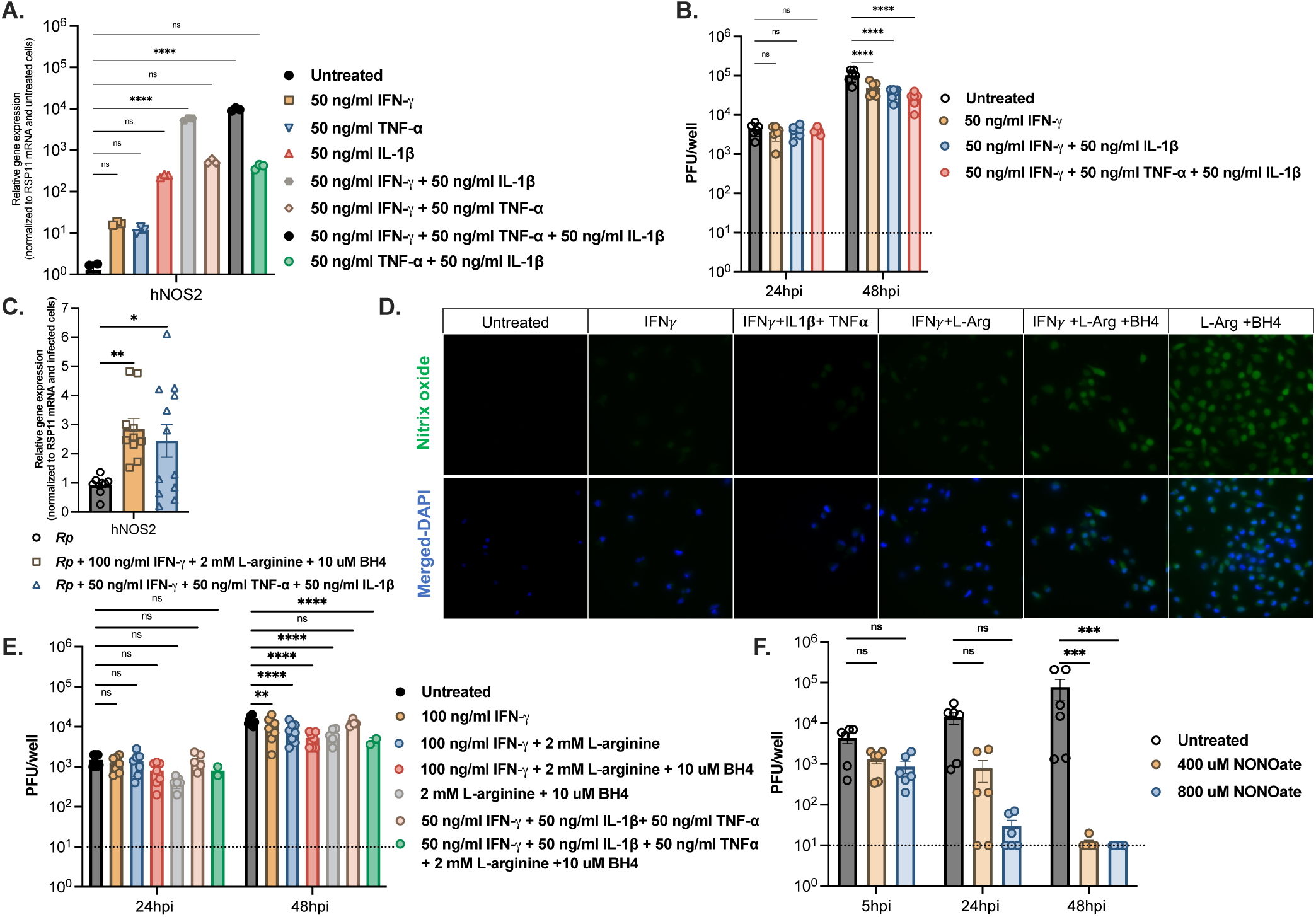
Human cells can produce iNOS and NO but only mildly restrict *R. parkeri*. **A** Human *Nos2* transcripts in A549s 6 h after cytokine treatment were quantified by qPCR and normalized to the housekeeping RSP11 gene. Data were then divided by untreated cells to reveal the fold change. Data are representative of two independent experiments, each consisting of three biological replicates. **B.** *R. parkeri* abundance in A549s, MOI of 0.1. The indicated amounts of IFN-γ, IL-1β, and TNF-α were pretreated for 6 h before infection; then they were added at 10 mpi and 24 hpi. Data are the combination of 3 independent experiments with 2 biological replicates each. **C.** Human *Nos2* transcripts of cytokine-treated and infected human MDMs at 48 hpi. Transcripts were measured by qPCR and normalized to the housekeeping gene RSP11. This value was then divided by infected/untreated cells to reveal the fold change. Human MDMs were derived from 4 healthy donors with n= 9, 10,12 biological replicates. **D**. DAF-2A staining of NO produced in cytokine-treated and infected human MDMs at 48 hpi, MOI of 0.1. Images are representative of two independent experiments using blood from two healthy donors. **E***. R. parkeri* abundance in human MDMs at 24 and 48 hpi, MOI of 0.1. The indicated amount of IFN-γ, IL-1*β*, TNF-*α*, L-arginine, and BH4 were added at 10 mpi and 24 hpi. Data are the combination of 3 independent experiments from 3 independent blood donors. At 24 hpi n= 7, 7, 7, 7, 6, 5, and 2 biological replicates and at 48 hpi n=8, 8, 8, 8, 6, 5, and 2 biological replicates. **F***. R. parkeri* abundance in human MDMs, MOI of 0.1. NONOate was added at 10 mpi and 24 hpi. Human MDMs were derived from n= 3 healthy donors and each experiment has 2 biological replicates. Asterisks indicate statistically significant differences. Statistics used a one-way ANOVA for panels A and C and a two-way ANOVA with Dunnett’s multiple comparisons test for panels B, E-G: *, p < 0.05; **, p < 0.01; ***, p < 0.001; ****, p < 0.0001.

iNOS requires the cofactor BH4 and the substrate L-arginine to produce NO (Gonçalves et al., 2021; Tayeh & Marletta, 1989). We hypothesized that human macrophage cells require additional BH4 and L-arginine to restrict *R. parkeri.* To determine whether the combination of cytokines plus cofactors increased NO, we performed immunofluorescence microscopy with DAF-2A, a fluorescent marker for NO. This revealed that NO was produced most robustly upon L-arginine+BH4 treatment, and that IFN-γ had little effect in increasing NO production either alone or with L-arginine + BH4 (**Fig. 3D**). To determine if this increase in NO correlated with increased killing of *R. parkeri,* we also measured PFUs over time in the presence of L-arginine and BH4. Supplementing the media of infected MDMs with BH4 + L-arginine only led to a 3-fold restriction of *R. parkeri.* When IFN-γ, TNF-α and IL-1β treatment was combined with excess L-arginine + BH4 during infection, the L-arginine + BH4 addition only minorly increased the restriction, from 1.1 to 3.3-fold, even upon adding these factors at two separate times postinfection (**Fig. 3E**). These data suggest that cytokines and cofactors can increase the amount of NO produced in human cells, but it does not meet the threshold to potently kill the bacteria.

The data suggested that perhaps human cells were unable to control *R. parkeri* even in the presence iNOS induction and iNOS-activating factors. We therefore hypothesized that NO was not sufficient to kill *R. parkeri* in human cells, perhaps due to the requirement of additional anti-rickettsial ISGs. To test this, we measured the ability of human cells to kill *R. parkeri* upon treatment with the NO donor NONOate. High doses (800 mM) of NONOATE strongly restricted *R. parkeri* in MDMs (1,000-fold, **Fig. 3F**), suggesting that in fact NO is sufficient to kill *R. parkeri* in human cells, but that human cells do not make such high concentrations under the conditions we tested. Together, these data suggest that cytokine cocktails and iNOS cofactors or substrates can induce and activate iNOS to make NO in human cells, but to they do not meet the threshold to restrict *R. parkeri*.

### *Nos2^-/-^* mice develop mild eschars upon *R. parkeri* infection

Eschars, also known as tache noire, are the hallmark human disease manifestation of *R. parkeri, R. conorii, O. tsutsugamushi* and other arthropod-borne pathogens and serve as key clinical feature for gross diagnosis of disease (Walker, 2003; Walker et al., 1988). In humans, eschars form within 5 days at the tick bite site and may take several weeks to heal completely (Burke et al., 2021; C. D. Paddock et al., 2008; Paddock et al., 2004). We previously reported that unlike WT mice, mice lacking both IFN-I and IFN-γ receptors form eschars at the intradermal site of infection and develop disseminated diseases characterized by weight loss and sometimes lethality (Burke et al., 2021). However, this mouse model is not an ideal mimic of human infection, as *R. parkeri* does not cause lethality in humans and humans have functional IFNAR and IFNGR. Our data showed that *Nos2* is a major anti-rickettsial gene *in vitro* induced by both IFN-γ (**Fig. 2**) and IFN-I (Burke et al., 2020), and we therefore hypothesized that NO controlled infection in mice.

To determine the anti-rickettsial role of NO *in vivo*, we used NONOATE and L-NIL treatment to determine if these NO chemical donors and inhibitors can rescue and suppress the eschar formation in *Ifnar^-/-^ Ifngr^-/-^* and WT mice, respectively. After intradermal infection, 6 consecutive days of 1 mg/kg NONOate treatment mildly but significantly reduced eschar severity (**Fig. 4A**). NONOate treatment caused no weight change between untreated and treated groups of *Ifnar^-/-^ Ifngr^-/-^* mice, suggesting that it did not dramatically reduce systemic disease (**Fig. 4A**). To determine if chemically blocking iNOS would increase disease in WT mice, we treated WT mice for 3 consecutive days with 15 mg/kg L-NIL. Interestingly, whereas vehicle-treated mice had no lesions, L-NIL-treated mice developed a mild but significantly more severe eschar at the intradermal site of infection (**Fig. 4B**). L-NIL treatment did not alter the weight between control and treated groups, suggesting that the systemic disease severity was mild (**Fig. 4B**). Thus, chemically inducing or inhibiting NO suggested an anti-rickettsial role for iNOS *in vivo*.

**Figure 4:**
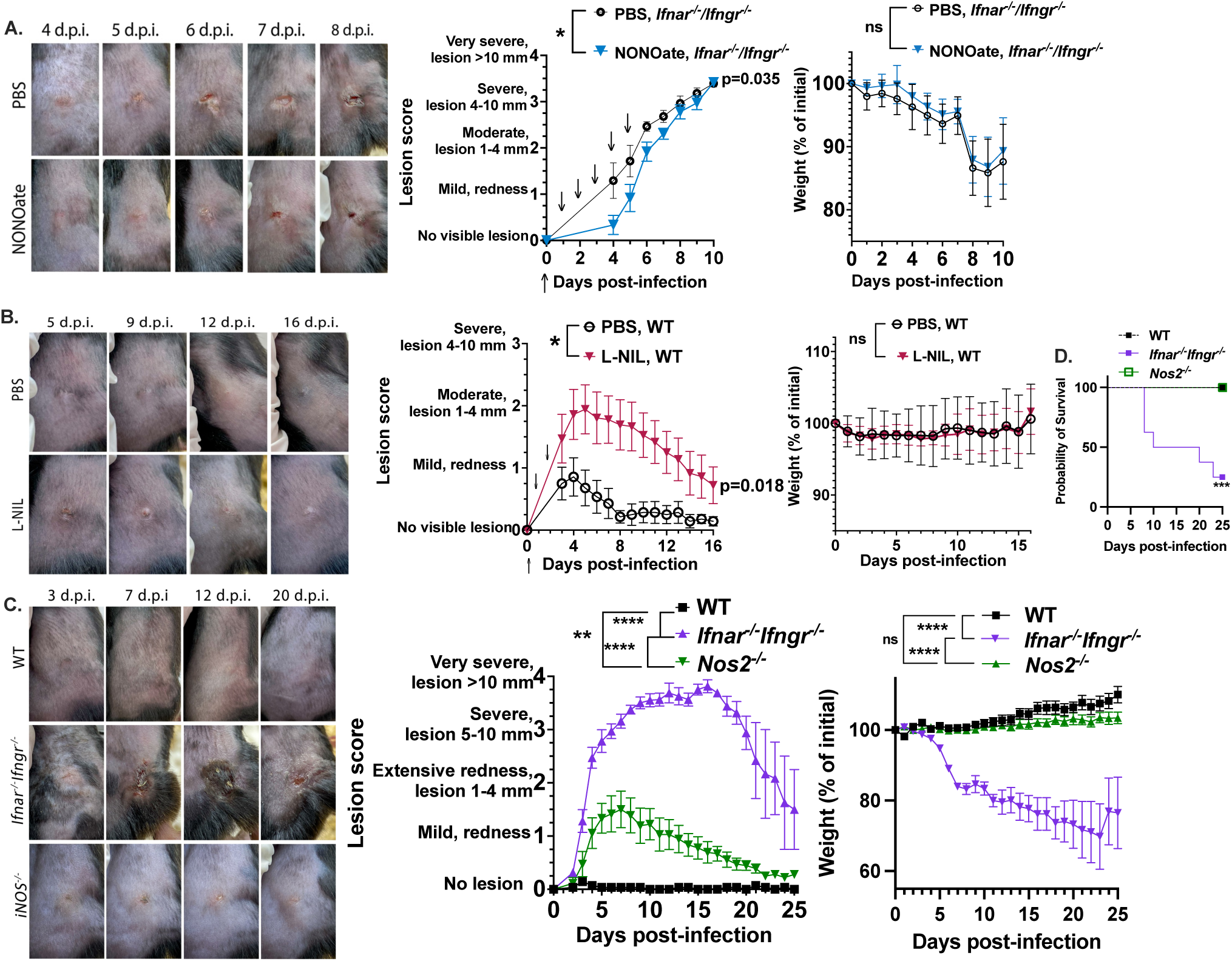
*Nos2*^-/-^ mice develop mild eschars and limited systemic disease upon *R. parkeri* skin infection. **A.** Representative images of *Ifnar^-/-^Ifngr^-/-^* mice intradermally infected on the right flank with 10^3^ PFU of *R. parkeri* and subcutaneously treated with 1 mg/kg NONOate (n=9 mice) and PBS (n=7 mice). The middle panel shows quantified lesion scores of mouse groups in the left pictures. Arrows indicate days of subcutaneous NONOate treatment. The right panel shows the weight changes of each group during the experiment. Data are the combination of two independent experiments. **B.** Representative images of WT mice intradermally infected with 5×10^6^ PFU *R. parkeri* and subcutaneously treated with 15 mg/kg of L-NIL (n=9 mice) or PBS (n=7 mice). The middle panel shows the quantified eschar lesion scores of mouse groups in the left pictures. Arrows indicate days of subcutaneous L-NIL treatment. The left panel shows the weight change of the mouse groups during infection. Data are the combination of two independent experiments. **C.** Representative images of intradermally infected skin from WT (n=7*), Ifnar^-/-^Ifngr^-/-^* (n=8), and *Nos2*^-/-^ (n=9) mice with 10^7^ PFU of *R. parkeri*. The graphs show the quantified lesion scores, weight changes, and **D.** survival. Data are combined from two independent experiments. All error bars indicate the standard error of the mean (SEM). The last mouse weight for deceased or euthanized *Ifnar^-/-^Ifngr^-/-^* mice was included in the statistical analysis until the last recorded day of the experiment. Statistics used two-way ANOVA for all panels of lesion scores or weight changes and Mantel-Cox for survival: *, p < 0.05; **, p < 0.01; ***, p < 0.001; ****, p < 0.0001.

We next asked whether infected *Nos2^-/-^* mice had increased disease as compared to WT mice, and whether these mice would serve as a non-lethal model of eschar-associated rickettsiosis. Intradermal infection of *Nos2^-/-^*mice caused mild but significantly more severe eschars as compared to infected WT control mice (**Figure 4C**). *Nos2^-/-^* mice developed eschars 4 days post-infection that were not as severe as in *Ifnar^-/-^Ifngr^-/-^* mice (**Fig. 4C**). *Nos2^-/-^* mice did not undergo significant weight loss, in contrast to *Ifnar^-/-^Ifngr^-/-^*mice, and there was no lethality in *Nos2^-/-^* mice (**Fig. 4C, 4D**), mimicking the non-lethal disease seen in humans (Arboleda et al., 2020). These mice therefore provide an additional animal model to study *R. parkeri* pathogenesis and reveal a key role for iNOS as an anti-rickettsial factor in rodents but not human cells.

## Discussion

It is often unclear why pathogens that circulate in rodent reservoirs, which are tolerant asymptomatic carriers, cause severe disease in humans. In this study, we identified *Nos2* as the major factor downstream of IFN-γ in mouse macrophages but not human macrophages that restricts the tick-borne human pathogen *R. parkeri*. Although iNOS is conserved from mice to humans and can produce NO, upregulating and activating iNOS in human cells was not sufficient to kill *R. parkeri*, suggesting that there is a threshold amount of NO needed to restrict *R. parkeri*, which can be met by mouse but not human cells. We leveraged this knowledge to show that *Nos2^-/-^* mice can serve as a non-lethal model to study eschar-associated rickettsiosis. Broadly, these findings suggest that differences in NO production between humans and rodents may account for susceptibilities to arthropod-borne pathogens that propagate themselves in wildlife reservoirs. Through the lens of evolution, this snapshot suggests that as humans interact less and less with wildlife, we may lose defenses to pathogens that are propagated in nature. Better understanding this phenomenon will have key implications for developing host-augmenting therapeutics and improved animal models to study infection.

A notable aspect of this study is the discrepancy between human and mouse iNOS in restricting *R. parkeri.* We found low *Nos2* upregulation in primary human cells during rickettsial infection in the presence of IFN-γ, that *Nos2* was induced by cytokine cocktails, and that NO production in human cells was increased by L-arginine and BH4 supplementation. Surprisingly, however, neither *Nos2* upregulation nor enhanced activity with L-arginine and BH4 dramatically reduced *R. parkeri* burdens. Chemical NO donors were sufficient to cause potent restriction, suggesting that NO itself is sufficient for the killing in human cells. These findings demonstrate that human iNOS is defective in both abundance and activity for restricting *R. parkeri*. Many explanations account for the differences of rodent versus human *Nos2/*iNOS regulation and activity, including transcriptionally, post-transcriptionally, and post-translationally (Azenabor et al., 2009; Blond et al., 2000; Bogdan, 2001a, 2001b; Panaro et al., 2003). *Nos2* gene expression is reduced by epigenetic gene silencing by CpG methylation, histone modifications, and chromatin compaction (Bogdan, 2015; Gross et al., 2014). Transcriptionally, IFN-γ alone does not upregulate human iNOS, but it can be upregulated by cytokine cocktails (Barilli et al., 2023; Kwon et al., 2001; Nussler et al., 1992; Zamora et al., 2000). Post-transcriptionally, microRNAs (miRNAs) such as miRNA-939 and miR-149 reduce iNOS protein abundance in human hepatocytes (Guo et al., 2012) and endothelial cells (Palmieri et al., 2014). Accordingly, little iNOS activity is found in cellular subfractions of human macrophages (SCHOEDON et al., 1993; Vodovotz et al., 1994; Weinberg et al., 1995). Our direct comparisons between mouse and human macrophages demonstrates that these regulatory discrepancies of iNOS result in important differences in susceptibility to infection.

Our findings that mouse iNOS restricts *R. parkeri* aligns with susceptibility of many other pathogens to NO in mouse cells. For example, with the obligate intracellular eukaryotic parasite *Toxoplasma gondii*, iNOS in mouse cells causes nitration of the pathogen-containing vacuoles and collapse of the intravacuolar membrane network (Zhao et al., 2024). In *Mycobacterium tuberculosis,* the production of NO and other reactive nitrogen intermediates (RNI) by macrophages helps to control infection (Darwin et al., 2003). In another example, NO produced by iNOS kills *R. rickettsii* in mouse macrophages upon IFN-γ stimulation, and in this study the authors suggest that NO kills *R. rickettsii* directly and that killing can be partially rescued by supplementing ATP, suggesting a role for NO in inhibiting metabolic processes (Fitzsimmons et al., 2021). However, while mouse iNOS can kill a variety of pathogens, the ability of human iNOS to restrict pathogens including *Rickettsia* was unclear. Our study highlights the importance of comparing both human and rodent cells when describing differences in innate immune-mediated restriction.

Our findings shed light onto how human versus rodent evolution shapes susceptibility to arthropod-borne pathogens. Ticks feed on a variety of animals worldwide, including birds, reptiles, amphibians, and a variety of mammals including the diverse and omnipresent Rodentia taxa. *R. parkeri* has been found in cattle, sheep, goats, small mammals like marsh rice rats, white-footed mice, house mice, meadow voles, black rats and wildlife such as black bears, feral pigs, white-tailed deer, cotton rats, and coyotes (Cumbie et al., 2020). Additionally, other *Rickettsia* can be found in opossums, capybaras, even some bats, birds, reptiles, amphibians and marsupials (Milagres et al., 2010; Santos-Silva et al., 2023; Seidi et al., 2024). Other arthropod-borne pathogens such as the mite-borne *Orientia tsutsugamushi,* which causes scrub typhus, are found in rodents, such as field mice, rats, and hamsters, while the Lyme disease agent *Borrelia burgdorferi* uses deer mice *Peromyscus* species as reservoirs (Barbour et al., 2023). Yet, over the past tens of thousands of years, humans have become less and less intertwined with wildlife. We speculate that over the course of *Homo sapiens* evolution, humans lost restrictive ISGs that are maintained in animals, which more routinely encounter tick-borne pathogens, and that iNOS represents such an example. Identifying additional antimicrobial ISGs that have been lost in humans will better elucidate the molecular explanation for human disease.

Another notable finding in our work is revealing *Nos2^-/-^* mice as a non-lethal animal model to study rickettsiosis. Several animal models have been examined to study *R. parkeri* virulence *in vivo.* This includes: different inbred mice that are all highly resistant to infection (Grasperge, Wolfson, et al., 2012), susceptible *Ifnar^-/-^Ifngr^-/-^*mice (Burke et al., 2021), guinea pigs that develop fevers (Stokes et al., 2020), non-human primates that developed eschars (Banajee et al., 2015), and using very high doses (10^8^) of *R. parkeri* in WT mice which can elicit lethality, although these are likely physiologically irrelevant doses (Londoño et al., 2019). We have found the *Ifnar^-/-^Ifngr^-/-^*mouse to be a robust model to study eschar formation, disease kinetics, bacterial virulence genes, and protective immunity (Borgo et al., 2022; Burke et al., 2021; Engström et al., 2019). However, it is not a fully faithful model to mimic disease because humans have functional IFN receptors. Furthermore, this model can cause lethality with high doses, but there are no reported lethal cases of healthy patients with *R. parkeri* infection (Paddock et al., 2004). Therefore, an ideal model would be an *ISG*-deficient mouse that lacks the ISGs that are missing or malfunctional in humans, which develops eschars and mild systemic disease. We found that *Nos2^-/-^* mice develop mild eschars and no systemic disease, perhaps better mimicking certain aspects of human infection. However, we were surprised that the phenotypes in *Nos2^-/-^*mice were not more severe, given that iNOS fully accounted for IFN-γ-mediated restriction of *R. parkeri* in BMDMs. A somewhat similar phenotype occurs during *Chlamydia* infection, as iNOS accounts for most of the IFN-γ-mediated killing *in vitro*, but *Nos2^-/-^* mice are mostly identical to WT and do not develop disease (Ramsey et al., 2001). The intracellular complement-like protein Plac8 somewhat complements iNOS-mediated killing of *C. muridaum* in mice (Johnson et al., 2012); however, the role for Plac8 in *Rickettsia* infection has not been explored. Future studies into this and into other maladapted human ISGs are needed to better explain the molecular basis for human disease and further improve the models to study *R. parkeri* and other pathogens.

Augmenting host ROS and NO may be a potential pan-antimicrobial approach, however an important aspect of our findings is revealing the challenges of augmenting human innate immunity to combat infection. NO donors have been shown in preclinical studies to suppress viruses, bacteria, protozoa and fungi *in vitro* and *in vivo*. In addition, phase II trials found that NO donors can reduce microbial infection (Bath et al., 2021). We found that inhibiting iNOS or using NO donors alters disease; however, the effects with these drugs were mild. Along with the *in vitro* findings that a certain high threshold of NO must be produced to restrict infection, it is likely that other strategies will be needed to dramatically unleash human iNOS as a therapeutic strategy. Further studies into augmenting iNOS and potentially other host ISG responses may lead to additional, broad-scale approaches to treat infectious disease.

## Material & Methods

### Bacterial isolation

*R. parkeri* strain Portsmouth was originally obtained from Dr. C. Paddock (CDC). Bacteria were expanded in confluent T175 flasks of African green monkey kidney epithelial Vero cells in DMEM 2% FBS. Each flask of Vero cells was infected with 5 x 10^6^ *R. parkeri*. At 5-7 days post-infection, infected cells were scraped and centrifuged at 3000 x rpm for 5min at 4 °C. Pelleted cells were then resuspended in K-36 buffer (0.05 M KH_2_PO_4_, 0.05 M K_2_HPO_4_, 100 mM KCl, 15 mM NaCl, pH 7) and dounced (60 strokes) on ice. The solution was centrifuged at 200xg for 2 min to pellet host cell debris. Supernatant containing *R. parkeri* was overlaid on Ficoll-Paque Plus (Cytiva) solution. Gradients were centrifuged at 20000 × rpm in Sorvall RC 5C for 20 min at 4 °C to obtain bacterial pellets. The bacterial pellets were resuspended in brain heart infusion (BHI) media for 200 ul per flask and stored at - 80°C. Bacterial PFUs were determined via plaque assays by serially diluting the bacteria in 12-well plates containing a monolayer of Vero cells. Prepared plates were then spun at 300xg for 5 min. At 24 hpi, the media for each well was replaced with 2 ml per well DMEM 5% FBS and 2.4% Avicel. At 7 dpi, 2 ml of 7% PFA was added to each well and incubated for 30 min before staining with 1.25% crystal violet solution for 15 min.

### Deriving BMDMs

Femurs, tibias and fibulas were excised from at least 6-week-old mice to obtain bone marrow. Connective tissue was removed from these bones to be sterilized with 70% ethanol. The bones were washed with BMDM media (20% FBS, 1% sodium pyruvate, 0.1% β-mercaptoethanol, 10% conditioned supernatant from 3T3 fibroblasts, in Gibco DMEM containing glucose and 100 U/mL penicillin and streptomycin) and ground using a pestle in a mortar. Bone homogenate was passed through a 70-μm nylon Corning Falcon cell strainer (Thermo Fisher Scientific, 08-771-2) to remove particulates. Filtrates were centrifuged in an Eppendorf 5810R at 300xg for 8 min, the supernatant was aspirated, and the remaining pellet was resuspended in BMDM media. Cells were then plated in non-tissue culture-treated 150 cm petri dishes (ratio of 10 dishes per 2 femurs/tibias) in 30 ml BMDM media and incubated at 37 °C. An additional 30 ml was added 3 days later. At day 7, the media was aspirated, and cells were incubated at 4 °C with 15 ml cold PBS (Gibco, 10010-023) for 10 min. BMDMs were then scraped from the plate, collected in a 50-ml conical tube and centrifuged at 300xg for 5 min. The PBS was then aspirated, and cells were resuspended in BMDM media with 10% FBS and 10% DMSO to be stored in liquid nitrogen.

### Deriving PBMCs

100 mL of different healthy donors was obtained from the UCI Blood Donation Center (Irvine, CA). Peripheral blood mononuclear cells (PBMCs) were isolated from buffy coats by Ficoll-Paque (Cyvita) gradient centrifugation after 30 minutes of 400xg centrifugation without brake. CD14+ monocytes were magnetically isolated from PBMCs with CD14+ coated beads (Miltenyi Biotec). Monocytes were cultured in a non-TC 100 cm culture dishes in Dulbecco’s modified Eagle’s medium (DMEM) with 10 ng/mL recombinant human-MCSF or GMCSF (R&D), 2 mM glutamine, 25 mM HEPES, 10% fetal bovine serum (FBS) and 5% human AB serum (103012-084 Access cell culture) grown at 37 °C with 5% CO2 for 5-7 days. Macrophages were physically scraped from the culture dish after incubating in cold phosphate-buffered saline (PBS) for 10 minutes at 4 °C. The cells were counted and plated onto TC-treated culture plates to prepare for infection.

### Infections in vitro

To plate cells for infection, ∼33,300 cells were plated into 96-well plates. Approximately 16 h later, a 30% preparation of *R. parkeri* was thawed on ice and diluted into fresh media to the desired MOI and concentration. Media was then aspirated from each well and replaced with 84 µl media containing *R. parkeri*; the plates were spun at 300xg for 5 min in an Eppendorf 5810R. Infected cells were then incubated at 33 °C and 5% CO2 for the duration of the experiment. For treatments with recombinant mouse IFNs (R&D) were added directly to infected cells immediately after spinfection. Human 100 ng/mL human IFN-γ (Peprotech), 2 mM L-Arginine (MP Biomedicals), 10 uM BH4 (MedChem), 50 ng/mL TNF-α or IL-1β (Peprotech) were added at 0 hpi and 24 hpi for human MDMs. Human IFN-γ, TNF-α, and IL-1β at 50 ng/mL doses were added to pre-treat A549 for 6 hours before infection and added back at 0 hpi and 24 hpi. DETA-NONOate (Cayman) was dosed with 400 uM and 800 uM concentrations at 0 hpi and 24 hpi. 1 mM L-NIL (Cayman) was only added right after spinfection for BMDMs.

To measure the PFU, at 5, 24, 48, and 72 hpi, supernatants from infected cells were aspirated from individual wells, and each well was gently washed twice with 168 µl sterile Milli-Q-grade water. 168ul sterile Milli-Q water was then added to each well and repeatedly pipetted up and down to lyse the host cells. Serial dilutions of lysates were added to confluent Vero cells in 12-well plates that were plated 24 or 48 h previously. Plates were then spun at 300g using an Eppendorf 5810R centrifuge for 5 min at room temperature and incubated at 33 °C overnight. At 24 hpi, media for each well were replaced with 2 ml per well DMEM 5% FBS and 2.4% Avicel. At 7 dpi, 2 ml of 7% PFA was added to each well and incubated for 30 min before staining with 1.25% crystal violet solution for 15 min before washing with generous water to quantify.

### RNA-sequencing

5 × 10^5^ BMDMs and human MDMs were plated in 24-well plates, infected with *R. parkeri* at MOI 2, and treated with 0.2 ng/mL of recombinant mouse or human IFN-γ. At 12 hpi, the cells were lysed, and RNA was purified using a RNeasy purification kit (Qiagen). RNA quality was assessed using an Agilent 2100 Bioanalyzer, and all samples had RNA integrity number (RIN) values above 8.0. Transcripts were selected using polyA selection (using Dynabeads mRNA Purification Kit, Invitrogen) and enzymatically fragmented using the Apollo library prep kits (Wafergen PrepX RNA library prep for Illumina). Libraries were constructed using Apollo 324 (IntegenX), PCR amplified and multiplexed. The resulting libraries were sequenced at the UCI Genomics Research& Technology Hub using single-end reads, 50-base length, with the Hiseq Illumina platform. Reads were aligned to the Mus musculus C57BL/6 reference genome (GRCm39) or human genome (GRCh38.p10) using STAR aligner version 2.7.10a. Reads were counted using the FeatureCounts command from the Subread package v2.0.3. Differential gene expression was performed using DESeq2.

### qPCR

Total RNA was isolated from induced A549 and human primary macrophages using the RNeasy mini kit (Qiagen). 1 μg of input RNA was used as a template for reverse transcription using ProtoScript II First Strand cDNA synthesis (NEB) and random primer mix. RT-qPCR was performed using 5 μl of 10-fold-diluted cDNA and primers targeting human *Nos2* (5’-TTC AGTATCACAACCTCAGCAAG-3’ and 5’-TGGACCTGCAAGTTAAAATCCC-3’) and RPS11 (5’-GCCGAGACTATCTGCACTAC-3’ and 5’-ATGTCCAGCCTCAGAACTTC-3’) with a SYBR green protocol of Luna Universal qPCR master mix (NEB) on the CFX96 Touch Real-time PCR Detection system (Biorad). qPCR conditions were as follows: initial denaturation step at 95 °C for 3 min, then 40 cycles of 95 °C for 10 sec, 59 °C for 60 sec. Transcript levels of *NOS2* induced by different cocktail treatments were determined by normalizing the target transcript C_T_ value to the C_T_ value of the endogenous housekeeping RPS11 transcript and uninfected or infected cells.

### DAF-2 DA staining

Human macrophages were treated with medium alone, IFN-γ, L-Arginine, BH4, or a combination of these as above and infected with WT *R. parkeri* (MOI of 0.1). Infected macrophages were incubated at 37°C with 5% CO2. After 48 h, the macrophages were stained with 4,5-diaminofluorescein diacetate (DAF-2 DA) (5 uM; Cayman Chemical) for 30 min at RT in the dark and fixed with PBS 4% paraformaldehyde (PFA) and stained with DAPI for 30 min. The slides were examined using a Leica DMi8 microscope.

### Mouse infections

Mice were between 8 and 35 weeks old at the time of initial infection. Mice selected for experiments were age and sex matched. The sex of mice used for survival after i.d. infection and raw data for mouse experiments are provided in the Source Data for each figure. Power analysis was performed to determine sample sizes before initial experiments. All mice were of the C57BL/6J background. All mice were healthy at the time of infection and were housed in microisolator cages and provided chow, water, and bedding. No mice were administered antibiotics or maintained on water with antibiotics. Experimental groups were littermates of the same sex that were randomly assigned to experimental groups. For mouse infections, *R. parkeri* was prepared by diluting 30%-prep bacteria into cold sterile PBS on ice, centrifuged to pellet the bacteria, and resuspended in PBS. Bacterial suspensions were kept on ice during injections. For i.d. infections, mice were anaesthetized with 2.5% isoflurane via inhalation. The right flank of each mouse was shaved with a hair trimmer (Braintree CLP-41590), wiped with 70% ethanol, and 50 µl of bacterial suspension in PBS was injected intradermally using a 29-gauge needle. Mice were monitored for ∼3 min until they were fully awake. No adverse effects were recorded from anesthesia. For subcutaneous injections, mice were scuffed, and a 27-gauge needle was used to deliver 50-100 µl of diluted L-NIL or NONOate within 1 cm of the infection site. All mice in this study were monitored daily for clinical signs of disease throughout infection, such as hunched posture, lethargy, scuffed fur, paralysis, facial edema, and lesions on the skin of the flank and tail. Mice were also monitored daily for changes in body weight. If a mouse displayed severe signs of infection, as defined by lethargy that prevented normal movement, body temperature below 90 degrees F, or a weight loss of >20% their starting weight, the animal was immediately and humanely euthanized using CO2 followed by cervical dislocation, according to IACUC-approved procedures. Pictures of skin lesions were obtained with permission from the Animal Care and Use Committee Chair and the Office of Laboratory and Animal Care. Pictures were captured with an Apple iPhone 11 Pro Max, software iOS 18.3.2, and were evenly exposed using Auto edit through Camera Raw Filter in Adobe Photoshop 2025.

## Acknowledgements

T.P.B. was supported by the National Institute of Allergy and Infectious Diseases (NIAID) of the National Institutes of Health (NIH) under Award Number R01AI185119. A.A.G. was supported by NIH-MARC U-STAR training grant T34GM136498. The content is solely the responsibility of the authors and does not necessarily represent the official views of the NIH. We thank Dr. Matthew Marsden at UC Irvine for assistance with CD14+ isolation protocols. We thank Dr. Orkide Koyuncu at UC Irvine for allowing us to use the Leica DMi8 microscope and Khanh Luong for guidance using this microscope. We thank the UC Irvine Genomic Core staff and Dr. Melanie Oakes for their help with the RNA library preparation and the sequencing service.

## Author contributions

A.P.L. performed *in vitro* experiments and designed *in vivo* experiments. A.A.G. performed *in vivo* experiments. N.D. performed the bioinformatic analysis of the RNA-sequencing. A.B. generated VSV and provided guidance on quantifying virus titer. A.P.L. wrote the original draft of this manuscript. T.P.B. contributed to experimental design, oversaw data management, the responsible conduct of research, and personnel, and critically reading and editing of this manuscript.

